# Population coding of valence in the basolateral amygdala

**DOI:** 10.1101/321232

**Authors:** Xian Zhang, Bo Li

**Author notes:** Correspondence to: Bo Li.

## Abstract

The basolateral amygdala (BLA) plays an important role in associative learning, by representing both conditioned stimuli (CSs) and unconditioned stimuli (USs) of positive and negative valences, and by forming associations between CSs and USs. However, how such associations are formed and updated during learning remains unclear. Here we show that associative learning driven by reward and punishment profoundly alters BLA neuronal responses at population levels, reducing noise correlations and transforming the representations of CSs to resemble the distinctive valence-specific representations of USs. This transformation is accompanied by the emergence of prevalent inhibitory CS and US responses, and by the plasticity of CS responses in individual BLA neurons. During reversal learning wherein the expected valences are reversed, BLA population CS representations are remapped onto ensembles representing the opposite valences and track the switching in valence-specific behavioral actions. Our results reveal how signals predictive of opposing valences in the BLA evolve during reward and punishment learning, and how these signals might be updated and used to guide flexible behaviors.

## Introduction

Many stimuli in the environment have valences. For example, food and water are attractive to animals and humans under metabolic demand, whereas harm and punishment are inherently aversive (Cardinal et al., 2002; Davis, 1998; LeDoux, 2000; Rosen, 2004; Schultz, 2006). Other environmental stimuli, such as a random sound or visual cue may not be attractive or aversive by nature. However, animals and humans have the ability to assign a positive or negative valence to an otherwise neutral stimulus (also known as conditioned stimulus, or CS) on condition that the stimulus is frequently associated with the occurrence of a good or bad consequence (also known as unconditioned stimulus, or US), and furthermore to revoke or reassign the valence if the stimulus-consequence contingency has changed (Lang and Davis, 2006; LeDoux, 2000; Pavlov, 1927; Schultz, 2006). This process, the core of which is known as associative learning, is fundamental for successfully foraging in a world filled with rewards, dangers and uncertainties, because it enables an organism to use, on the basis of past experiences, arbitrary environmental cues (CSs) to predict beneficial or detrimental outcomes (USs), and moreover to flexibly update the predictions in the face of changes in CS-US contingencies (Lang and Davis, 2006; LeDoux, 2000; Pavlov, 1927; Schultz, 2001, 2006).

How the brain assigns valences to CSs according to CS-US contingencies, and updates the valences when the contingencies change, has been a subject of intensive study. Substantial evidence indicates that neurons in the basolateral amygdala (BLA) have an important role in such associative learning (Duvarci and Pare, 2014; Grundemann and Luthi, 2015; Herry and Johansen, 2014; Janak and Tye, 2015; Johansen et al., 2011; Maren and Quirk, 2004). The BLA receives sensory inputs of all modalities, which can serve as CSs (Fendt and Fanselow, 1999; Gallagher and Holland, 1994; Gore et al., 2015; Janak and Tye, 2015; Lang and Davis, 2006; McDonald, 1998; Russchen et al., 1985; Sah et al., 2003; Salzman and Fusi, 2010; Sarter and Markowitsch, 1985), as well as inputs carrying appetitive or aversive information that may serve as USs (Belova et al., 2007; Bermudez and Schultz, 2010; Gore et al., 2015; Knapska et al., 2007; Livneh and Paz, 2012; Muramoto et al., 1993; Paton et al., 2006; Romanski et al., 1993; Shabel and Janak, 2009; Tye et al., 2008; Uwano et al., 1995; Wolff et al., 2014; Yu et al., 2017). Notably, recent studies suggest that BLA neurons (or at least some of them) responsive to appetitive and aversive USs – which are identified and targeted on the basis of expression of the immediate early gene *c-fos* or projection targets – are hard-wired in valence-specific circuits, as activation of these neurons optogenetically induces behavioral responses conforming to the valences of the actual USs, and moreover can substitute for the USs to drive appetitive or aversive conditioning (Gore et al., 2015; Kim et al., 2016; Namburi et al., 2015; Redondo et al., 2014; Stuber et al., 2011).

An emerging theme is that during associative learning CSs acquire the ability to activate the hard-wired US circuits, which in turn drives valence-specific behavioral responses (O’Neill et al., 2018). Consistent with this idea, CS-evoked spiking activity in individual BLA neurons increases with appetitive or aversive conditioning, correlates with learning and represents the valence of the US (Belova et al., 2007; Paton et al., 2006; Shabel and Janak, 2009). Notably, at single cell levels, these neurons update CS-US associations (and thus the valences of the CSs) when CS-US contingencies change (Paton et al., 2006). Nevertheless, these studies mainly focused on BLA neuronal responses in well-trained animals, leaving uncertain how these valence-specific responses develop during learning. Furthermore, the responses of individual neurons in single trials are noisy and thus do not reliably predict behavior (Renart and Machens, 2014). A recent study used imaging methods to simultaneously record the activities of ensembles of BLA neurons throughout fear conditioning, and showed that BLA population activities in single trials, extracted using dimensionality reduction methods, provide a robust account for learning-induced freezing behavior (Grewe et al., 2017). This approach provides an opportunity to examine the relationship between BLA population activities and the establishment of valence-specific behaviors across timescales, such as, for example, how population CS responses in the BLA evolve during learning to represent both positive and negative valences, and how these representations may be dynamically updated in response to changes in CS-US contingencies and thus influence ongoing behaviors.

In this study, we monitored the activities of ensembles of BLA neurons – reported with the genetically encoded calcium indicator GCaMP6f (Chen et al., 2013) – by imaging through gradient-index (GRIN) lenses (Ghosh et al., 2011; Resendez et al., 2016; Yu et al., 2017) implanted in the BLA in mice performing a Pavlovian associative learning task, in which mice learned to associate one CS with reward, and the other with punishment. We subsequently imaged the activities of these neurons in a reversal learning procedure in which the valences initially assigned to the CSs were reversed. Our results illustrate how signals predictive of opposing valences in the BLA develop during learning, and how these signals are updated on a trial-by-trial basis during reversal learning in a manner that they can be used to guide appropriate and flexible behavioral responses.

## Results

### The innate responses of BLA neurons to CSs and USs of opposing valences

To monitor neuronal activity in the BLA in behaving animals, we injected the BLA of wild type mice with an adeno-associated virus (AAV) expressing GCaMP6f (AAV1-Syn-GCaMP6f.WPRE.SV40) (Fig. 1a, b). The GCaMP6f delivered by this virus was predominantly expressed in excitatory pyramidal neurons (only 4.9±1% of GCaMP6f-expressing cells are GABAergic; n = 3 mice) (Supplementary Fig. 1a). We subsequently implanted a GRIN lens into the BLA and above the infected neurons (Fig. 1a, b; Supplementary Fig. 1b) in each of these mice. Four to six weeks after the surgery, we used a miniature integrated fluorescence microscope (Ghosh et al., 2011; Resendez et al., 2016) to record through the GRIN lenses dynamic GCaMP6f fluorescent signals from BLA neurons in these mice in wakefulness and under head-restrain (Methods). The constrained non-negative matrix factorization (CNMF) methods were used for imaging data processing before analysis, as previously described (Yu et al., 2017; Zhou et al., 2018) (Supplementary Video 1; Methods).

**Fig. 1.**
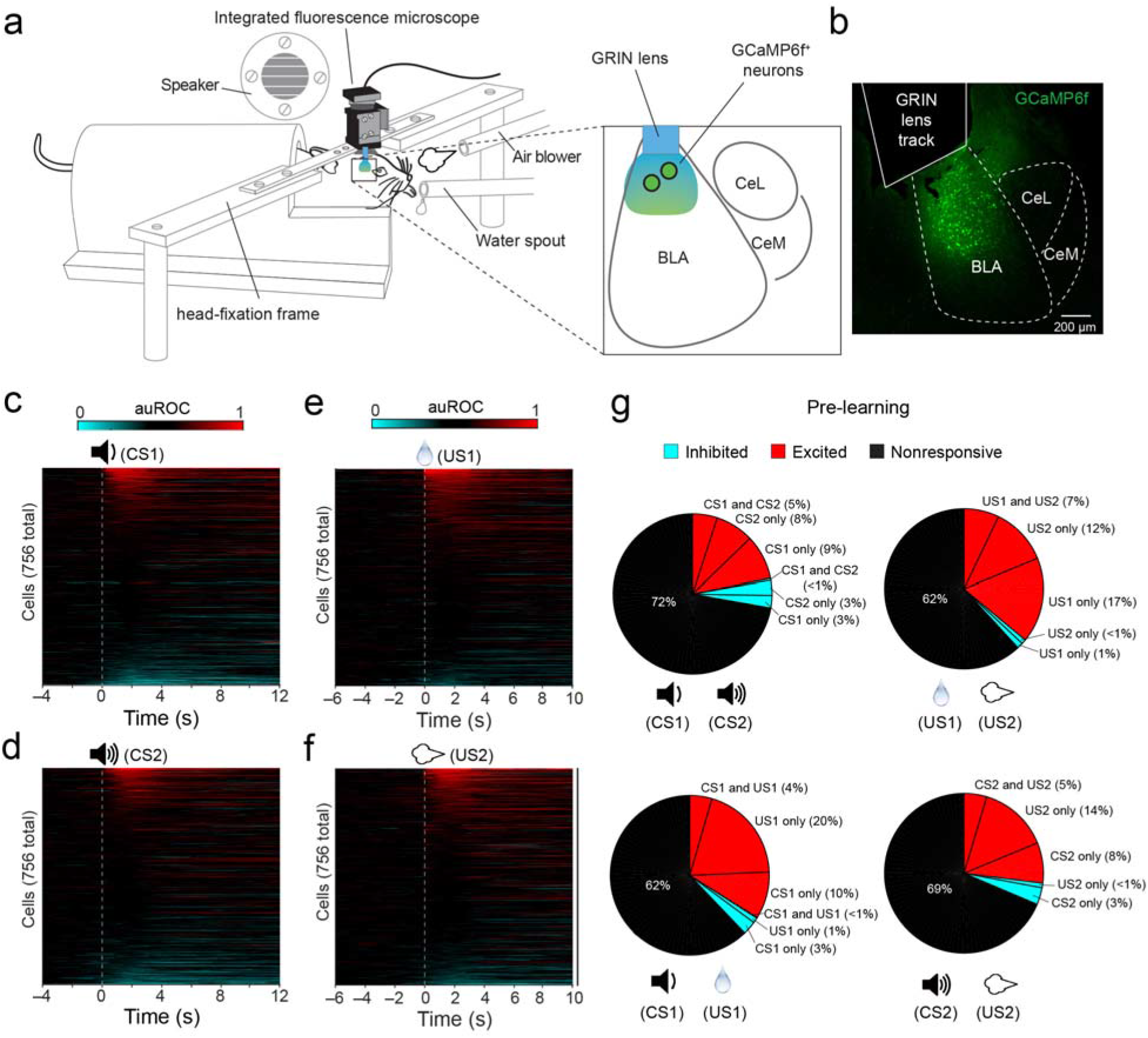
The innate responses of BLA neurons to CSs and USs. (a) A schematic of the setup for simultaneously monitoring behavioral and neuronal responses in head restrained mice. We imaged GCaMP6 signals in BLA neurons through GRIN lenses in behaving mice using a miniature integrated fluorescence microscope mounted on the head. (b) A representative confocal image of a coronal brain section containing the BLA, in which the track of an implanted GRIN lens was on top of the BLA neurons expressing GCaMP6f. (c) Heatmaps of auROC values for all neurons (n = 756 neurons, 6 mice) in trials in which CS1 was presented (indicated by the dashed line). Each row represents the temporal auROC values for one neuron, generated by comparing the calcium activity in this neuron at each time point (i.e., each picture frame) with that during baseline. Neurons are sorted according to their average auROC values during the 3-s time window after CS1 onset. (d-f) Same as (c), except that CS2 (d), US1 (water reward) (e), or US2 (airpuff) (f) was presented as the stimulus. (g) Pie charts showing the percent distributions of neurons responsive to different stimuli before learning. Note that at this stage, inhibitory responses were rare; and only a small percentage of neurons responded to both CS1 and US1, or to both CS2 and US2.

We first examined BLA neuronal activities in naïve mice in response to stimuli with different valences, including neutral tones (CS1s (2-kHz) and CS2s (10-kHz)), water rewards (US1s; note that mice were mildly water deprived) and aversive air-puffs blowing to the face (US2s) (Fig. 1a). BLA neurons exhibited diverse response profiles to these stimuli, with some responding to only one particular CS or US, while others responding to more than one stimuli (Supplementary Fig. 2). To characterize the dynamics of stimulus-evoked activities in each neuron, we performed receiver operating characteristic (ROC) analysis by comparing the GCaMP6f signals at baseline with those at different time points with respect to stimulus onset, and computed the values of the area under the ROC curve (auROC) (Supplementary Fig. 3; Methods). We used a permutation test to determine, for each neuron, whether the stimulus-evoked responses were statistically different (P < 0.05) from baseline (see Methods).

In naïve mice (n = 6), BLA neurons (n = 756) showed increased, decreased or no change in activity in response to a particular stimulus (Fig. 1c-f). In particular, about 14% and 13% of neurons were excited by CS1 and CS2, respectively, 24% and 19 % of neurons were excited by US1 and US2, respectively, while the majority of neurons showed only spontaneous activities (Fig. 1g). Only a small fraction of neurons was inhibited by any of these stimuli (<3%Fig. 1g).

There were also small fractions of neurons responsive to both CS1 and CS2 (5%), or to both US1 and US2 (7%) (Fig. 1g, top panels), and neurons responsive to both CS1 and US1 (4%), or to both CS2 and US2 (5%) (Fig. 1g, bottom panels).

It has been shown that different but partially overlapping populations of BLA neurons respond to reward and punishment (Gore et al., 2015; Kim et al., 2016; Namburi et al., 2015; Paton et al., 2006). Consistent with these studies, we found that among the US-responsive BLA neurons, 47.2% were excited by US1 (water reward) but not US2 (air-puff), and 33.3% were excited by US2 but not US1 (Supplementary Fig. 4; Fig. 1g, upper right). There was also a smaller population (19.4%) excited by both USs (Supplementary Fig. 4; Fig. 1g, upper right). This latter population may represent neurons responsive to salience (Belova et al., 2007). Notably, the neurons excited by the USs of opposing valences were distributed in the field of view with no obvious anatomical separation (Supplementary Fig. 4), a feature consistent with the findings in previous studies in which BLA neuronal activity was monitored electrophysiologically or based on the expression of the immediately early gene *c-fos* ((Gore et al., 2015; Paton et al., 2006); but see (Kim et al., 2016)).

### Both reward learning and punishment learning link CS and US representations in BLA neurons

After imaging the activities of BLA neurons in naïve mice, we went on to train these mice in Pavlovian associative learning tasks and examined how these neurons might participate in learning. We trained the mice to first associate CS1 with US1 (reward learning), and then CS2 with US2 (punishment learning) (Fig. 2a; Methods). As the training progressed, the mice increased their licking in response to CS1 presentations (Fig. 2b, d; Supplementary Video 2) and eye closing (or “blinking”) in response to CS2 presentations (Fig. 2c, e; Supplementary Fig. 5, and Supplementary Video 3). These anticipatory responses provided measures of learning driven by US of either positive or negative valence. All mice reached high performance levels within 7 sessions of training, licking the spout in anticipation of water delivery and blinking the eye in anticipation of air puffing after CS onset and during the trace interval before US onset in over 90% of the trials (Fig. 2d, e).

**Fig. 2.**
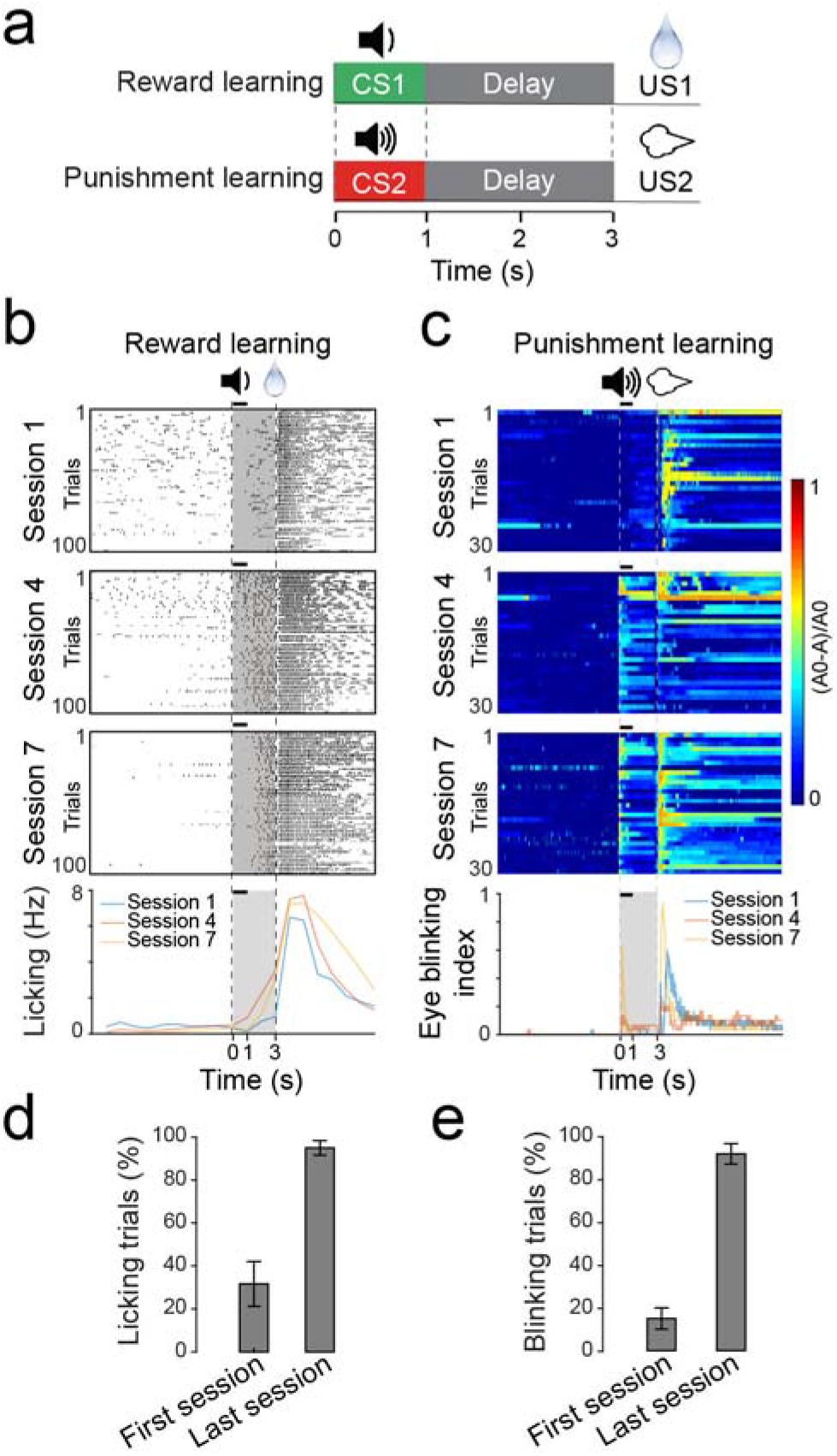
The behavioral task. (a) A schematic of the behavioral procedure, in which mice were trained to associate CS1 (a 2 kHz tone) with US1 (water reward), and CS2 (a 10 kHz tone) with US2 (airpuff blowing to the face). (b) Changes in licking behavior during the reward learning for a representative mouse. The upper three panels are raster plots of licking events during early (session 1), mid (session 4) and late (session 7) training stages. The bottom panel shows average licking rate over time (1-s bin) for each of the three sessions in the upper panels. Licks are aligned to the onset of CS1 (t = 0; the duration of CS1 (1 s) is indicated by a black bar above each panel). The delivery of US1 (water) was at 3 s after CS1 onset. Licks in the shaded area represent predictive licking events. (c) Eye blinking index, measured as (A0–A)/A0 (where A0 is the median eye size during the 10 s baseline, and A is the eye size in each picture frame), during the punishment learning for the same mouse as that in (b). The upper three panels are heat-maps of eye blinking during early (session 1), mid (session 4) and late (session 7) training stages. The bottom panel shows average blinking over time for each of the three sessions in the upper panels. Eye blinks are aligned to CS2 onset (t = 0; the duration of CS2 (1 s) is indicated by a black bar above each panel). The delivery of US2 (airpuffs) was at 3 s after CS2 onset. Eye blinks in the shaded area represent predictive blinking events. (d & e) The percentage of trials in which mice showed predictive licking (d) or blinking (e) (i.e., trials with at least one lick (d) or blink (e) event in the shaded area) in the first and last training sessions (the first 10 trials of each session were used for analysis) (n = 6 mice).

In the last training session (equal to or earlier than the 7^th^ session depending on the performance of individual mice) in the reward or punishment learning task, we imaged the activities of BLA neurons (n = 677) in these mice (n = 6) in response to CS1 followed by US1 (Fig. 3a) or CS2 followed by US2 (Fig. 3b), respectively. Compared with BLA neurons in naïve (pre-learning) mice, BLA neurons in the same but well trained (post-learning) mice showed interesting changes in the responses (Fig. 3c-e). First, the fraction of neurons that were excited by both CS1 and CS2 reduced from 5% (Fig. 1g, upper left) to 2% (Fig. 3C, upper left) (P = 0.0028, χ^2^ test), while the fraction of neurons excited by either CS1 or CS2 did not change (Fig. 3d), suggesting that learning reduces the overlap between CS representations (also see results in Fig. 7 below). Second, the fractions of neurons showing inhibitory responses to the CSs or USs were all markedly increased (Fig. 1g; Fig. 3c, e). Third, the fraction of neurons that were excited by both CS1 and US1 (Fig. 4a; also see the lower left panels of Fig. 1g and Fig. 3c), or by both CS2 and US2 (Fig. 4a; also see the lower right panels of Fig. 1g and Fig. 3c), was more than doubled, despite the fact that the fraction of neurons excited by either US1 or US2 alone was significantly reduced (Fig. 3d). The reduction in the number of US-excited neurons when the US was signaled by the CS is consistent with previous findings that expectation suppresses US responses of BLA neurons (Belova et al., 2007; Johansen et al., 2010). Fourth, the fraction of neurons that were inhibited by both CS1 and US1 (Fig. 4b; also see the lower left panels of Fig. 1g and Fig. 3c), or by both CS2 and US2 (Fig. 4b; also see the lower right panels of Fig. 1g and Fig. 3c), was also markedly increased.

**Fig. 3.**
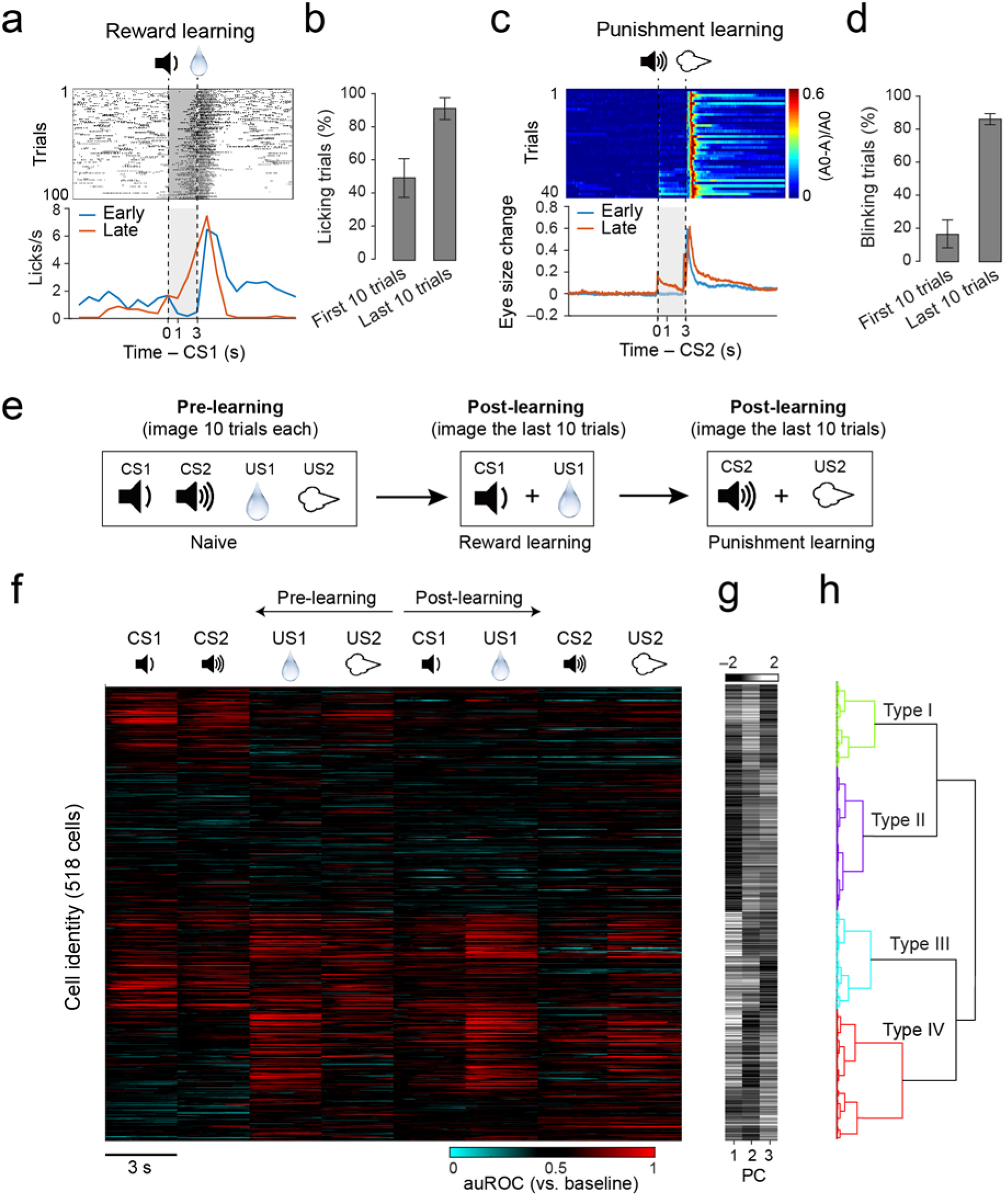
The CS and US responses in BLA neurons after learning. BLA neuronal activities were imaged in mice (n = 6) well trained with both the reward and the punishment conditioning. (a) Left: heatmaps of auROC values for all neurons (n = 677) in the reward block, in which CS1 was paired with US1 (indicated by the dashed lines). Each row represents the temporal auROC values for one neuron. Neurons are sorted according to their average auROC values during the 0–3 s time window. (b) Same as (a), except that imaging was performed in the punishment block, in which CS2 was paired with US2. (c) Pie charts showing the percent distributions of neurons responsive to different stimuli after learning. Note that at this stage, inhibitory responses became prominent; and the percentage of neurons responsive to both CS1 and US1, or to both CS2 and US2 increased. (d) Proportions of BLA neurons showing excitatory responses to the CSs and USs before and after learning (CS1, P = 0.09; CS2, P = 0.24; US1, ***P = 1.18e–07; US2, *P = 0.01; χ^2^ test). (e) Proportions of BLA neurons showing inhibitory responses to the CSs and USs before and after learning (CS1, ***P = 2.95e–09; CS2, ***P = 1.94e–12; US1, ***P = 1.78e–19; US2, ***P = 1.78e–19; χ^2^ test).

**Fig. 4.**
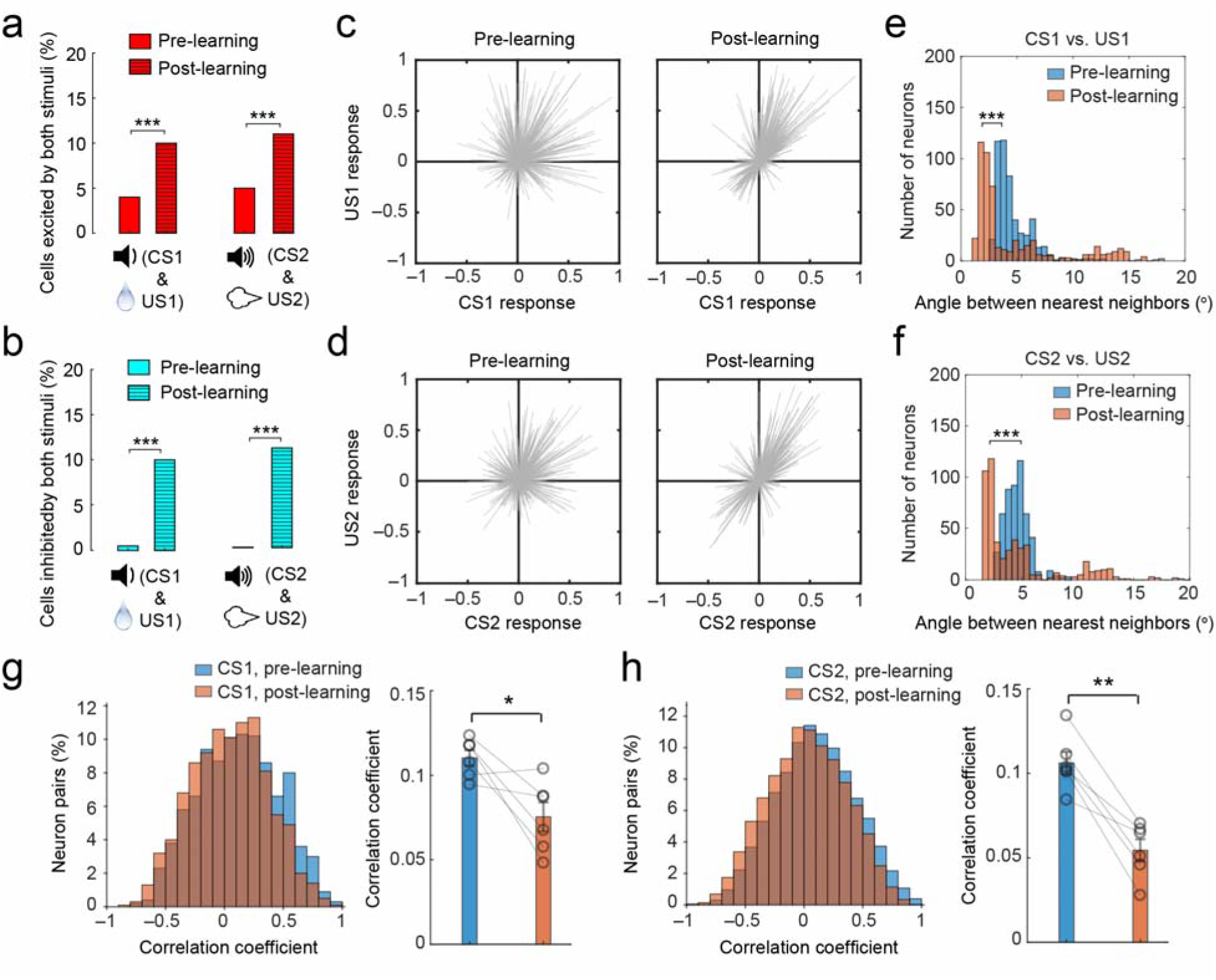
Learning links CS and US representations and reduces noise correlations in the BLA. (a) The percentage of BLA neurons showing excitatory responses to both CS and US (CS1 & US1, ***P = 5.3e–6; CS2 & US2, ***P = 3.2e–5; c^2^ test). (b) The percentage of BLA neurons showing inhibitory responses to both CS and US (CS1 & US1, ***P = 1.8e–16; CS2 & US2, ***P = 1.0e–20; c^2^ test). (c, d) The responses to CS1 and US1 (c), or to CS2 and US2 (d), for each neuron. Each line is a vector representing the responses of a particular neuron to both the CS and the US (values represent normalized auROC). Note that before learning, the vectors are distributed uniformly in the four quadrants, whereas after learning, the vectors are concentrated in quadrants I & III. (e) The distribution of angles between the nearest neighbors among vectors in (c) (pre-learning median, 4.07, n = 756 neurons, post-learning median, 2.68, n = 677 neurons, ***P = 4.25e–23, rank sum test). (f) The distribution of angles between the nearest neighbors among vectors in (d) (pre-learning median, 4.56, n = 756 neurons, post-learning median, 3.06, n = 677 neurons,***P = 1.18e–17, rank sum test). (g) Left, histograms of noise correlations in the responses to CS1 in BLA neurons in a representative mouse. Right, comparison of noise correlations between pre-learning and post-learning conditions (n = 6 mice, *p = 0.0285, paired t test). (h) Left, histograms of noise correlations in the responses to CS2 in BLA neurons in a representative mouse. Right, comparison of noise correlations between pre-learning and post-learning conditions (n = 6 mice, **p = 0.0015, paired t test). The bar graphs on the right in g & h represent mean ± s.e.m.

Overall, these changes suggest that learning substantially modifies the activity profile of BLA neurons. In particular, the third and fourth observations above (Fig. 4a, b) suggest that, after learning, a population of BLA neurons shows consistent responses to both a CS and the associated US. Supporting this notion, further analysis revealed that the vectors representing the CS and US responses of individual neurons were distributed randomly before learning, but were mainly confined to two opposing quadrants after either the reward or punishment learning (Fig. 4c-f), suggesting that the CS is linked up with and thus becomes predictive of the ensuing US with learning in an ensemble of BLA neurons. This finding is consistent with previous studies showing that CS and US responses are correlated after learning at the single cell level (Belova et al., 2007; Belova et al., 2008).

### Reward and punishment learning reduce noise correlations in the BLA

A critical feature of information encoding in neuronal ensembles is that noise correlations – the correlations between the responses of pairs of neurons to repeated presentations of an identical stimulus – affect the ability of downstream neurons to decode the information (Cohen and Kohn, 2011; Francis et al., 2018). Cognitive processes such as learning and attention can reduce noise correlations in cortical areas (Cohen and Kohn, 2011; Francis et al., 2018; Herrero et al., 2013; Mitchell et al., 2009; Miura et al., 2012; Ni et al., 2018; Shadlen and Newsome, 1998). To determine if such reduction occurs in the BLA, we computed the coefficients of noise correlations based on the CS responses of pairs of simultaneously recorded BLA neurons in each mouse (n = 6) (Methods), for both the pre-learning and the post-learning conditions (Fig. 4g, h). The noise correlations during CS1 presentations and those during CS2 presentations were both reduced after learning (Fig. 4g, h). These reductions could be important for the expression of the valence-specific behavioral responses, as a reduction in noise correlations is thought to increase the signal-to-noise of population responses and behavioral performance (Cohen and Kohn, 2011; Francis et al., 2018; Herrero et al., 2013; Mitchell et al., 2009; Miura et al., 2012; Ni et al., 2018; Shadlen and Newsome, 1998).

### Both reward and punishment transform CS representations in BLA neurons

Our results thus far indicate that both reward learning and punishment learning profoundly change BLA neurons’ responses to the CSs, which are presumably important for guiding appropriate behaviors. However, it is unclear how these changes evolve during learning. To address this issue, we took advantage of the mice (3 out of 6) that learned both the reward and the punishment tasks within one session of training (Fig. 5a-d), and thus allowed us to unambiguously track with imaging the activities of the same BLA neurons before and after learning (Fig. 5e) (see Methods).

**Fig. 5.**
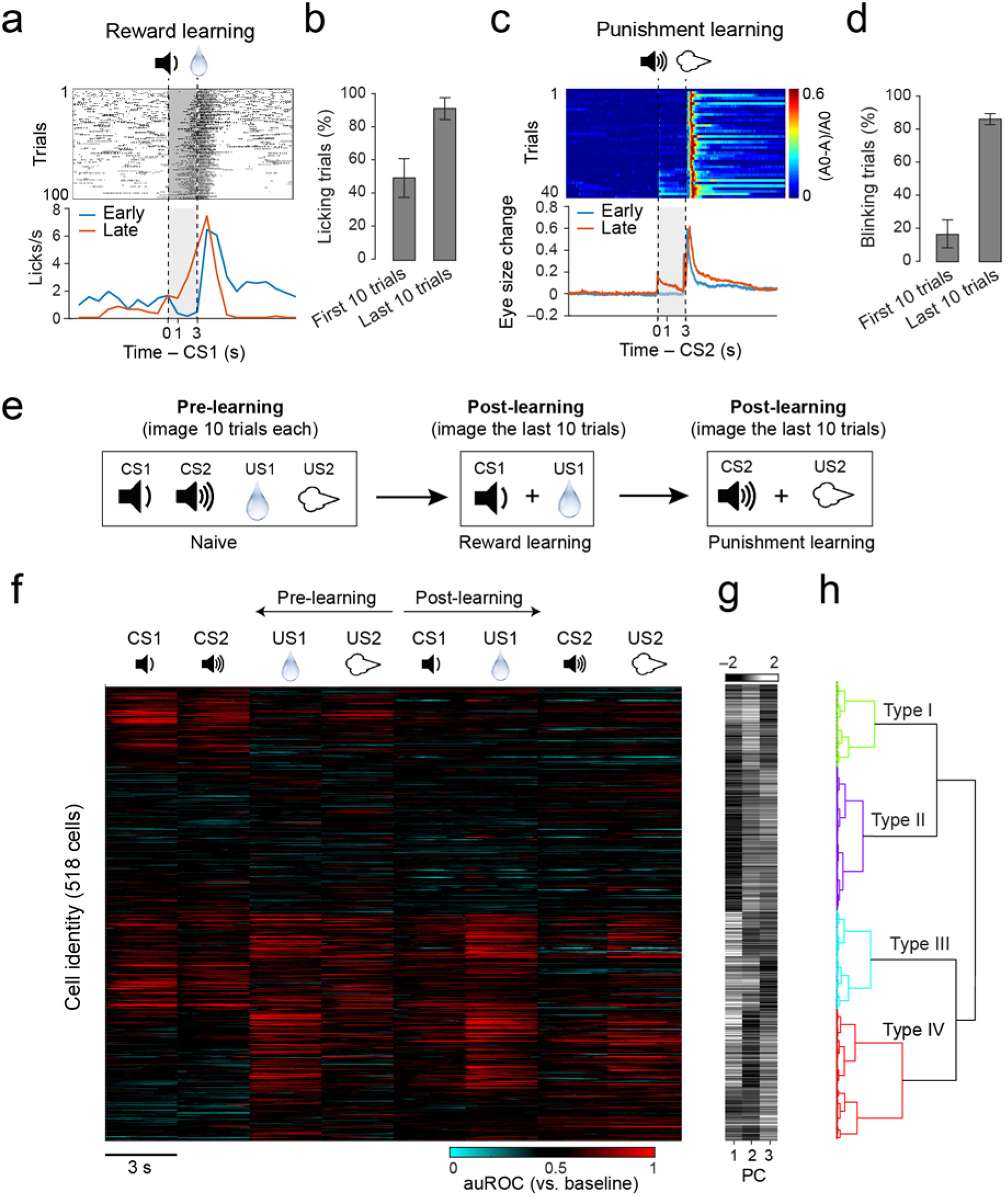
Tracking the activities of the same neurons in the BLA during learning. These mice (n = 3) learned both the reward and the punishment tasks within one session, thus eliminating the need for acquiring imaging data in different days (which inevitably causes shift in the neuronal populations being imaged, due to the detachment and reattachment of the camera across days), and allowing us to unambiguously track the same neurons before and after learning. (a) Changes in licking behavior during the reward learning for a representative mouse. The upper panel is a raster plot of licking events across trials. The bottom panel shows average licking rate over time (1-s bin), plotted separately for early and late trials. Licks are aligned to the onset of CS1 (t = 0; the duration of CS1 was 1 s). The delivery of US1 (water) was at 3 s after CS1 onset. Licks in the shaded area represent predictive licking events. (b) The percentage of trials in which mice showed predictive licking in the first and last 10 trials (n = 3 mice). (c) Changes in eye blinking behavior during the punishment learning for the same mouse as that in (a). In the upper panel are the heat-maps of eye size changes across trials. The bottom panel shows average blinking over time, plotted separately for early and late trials. Eye blinks are aligned to CS2 onset (t = 0; the duration of CS2 is 1 s). The delivery of US2 (airpuffs) was at 3 s after CS2 onset. Eye blinks in the shaded area represent predictive blinking events. (d) The percentage of trials in which mice showed predictive blinking in the first and last 10 trials (n = 3 mice). (e) The timing of the imaging experiments relative to behavioral training. (f-h) Classification of BLA neurons based on their response profiles across learning. The same neurons were unambiguously tracked before and after learning. (f) Heat-maps of stimulus-evoked responses, represented as auROC values, for individual neurons before and after learning. Each row represents one neuron (518 in total from 3 mice). (g) The first three principal components of the auROC profiles for each neuron. Each row represents the corresponding neuron in (f). (h) Hierarchy clustering of all neurons based on the PCA analysis.

As a first step to characterize how BLA neurons change their responses during learning, we classified these neurons (n = 518) into distinct functional types by hierarchical clustering following principal component analysis (PCA) on their responses to different stimuli, including CS1, CS2, US1 and US2 before and after learning in the reward and punishment learning tasks. This clustering yielded four types of neurons (Fig. 5f-h). Although there is heterogeneity within these types, the clustering captured some of their major features. Type I neurons (n = 97) on average had high excitatory responses to CS1 and CS2 before learning, had relatively low responses to USs (especially to US1), and markedly reduced their excitatory responses to or even became inhibited by the CSs after learning (Fig. 6a-c). Type II neurons (n = 162) on average had inhibitory responses or low excitatory responses to the CSs and USs, with the CS responses remaining largely stable during learning (Fig. 6d-f). Type III neurons (n = 112) on average had high excitatory responses to the CSs and USs before learning, and overall reduced their excitatory responses to the CSs after learning (Fig. 6g-i). Type IV neurons (n = 147) on average had inhibitory or low excitatory responses to CS1 and CS2 before learning, had relatively high excitatory responses to the USs (especially to US1), and many of them substantially increased their excitatory responses to the CSs after learning (Fig. 6j-l). The wide-spread depression and inhibition, and the more restricted potentiation of CS responses among BLA neurons after learning can, at least in part, account for the major effects of learning that we have observed, such as the marked increase in the fraction of neurons showing inhibitory response to the CSs (Fig. 1g; Fig. 3c,e), and the increase in the fraction of neurons showing excitatory response to both CS1 and US1, or to both CS2 and US2 (Fig. 4a).

**Fig. 6.**
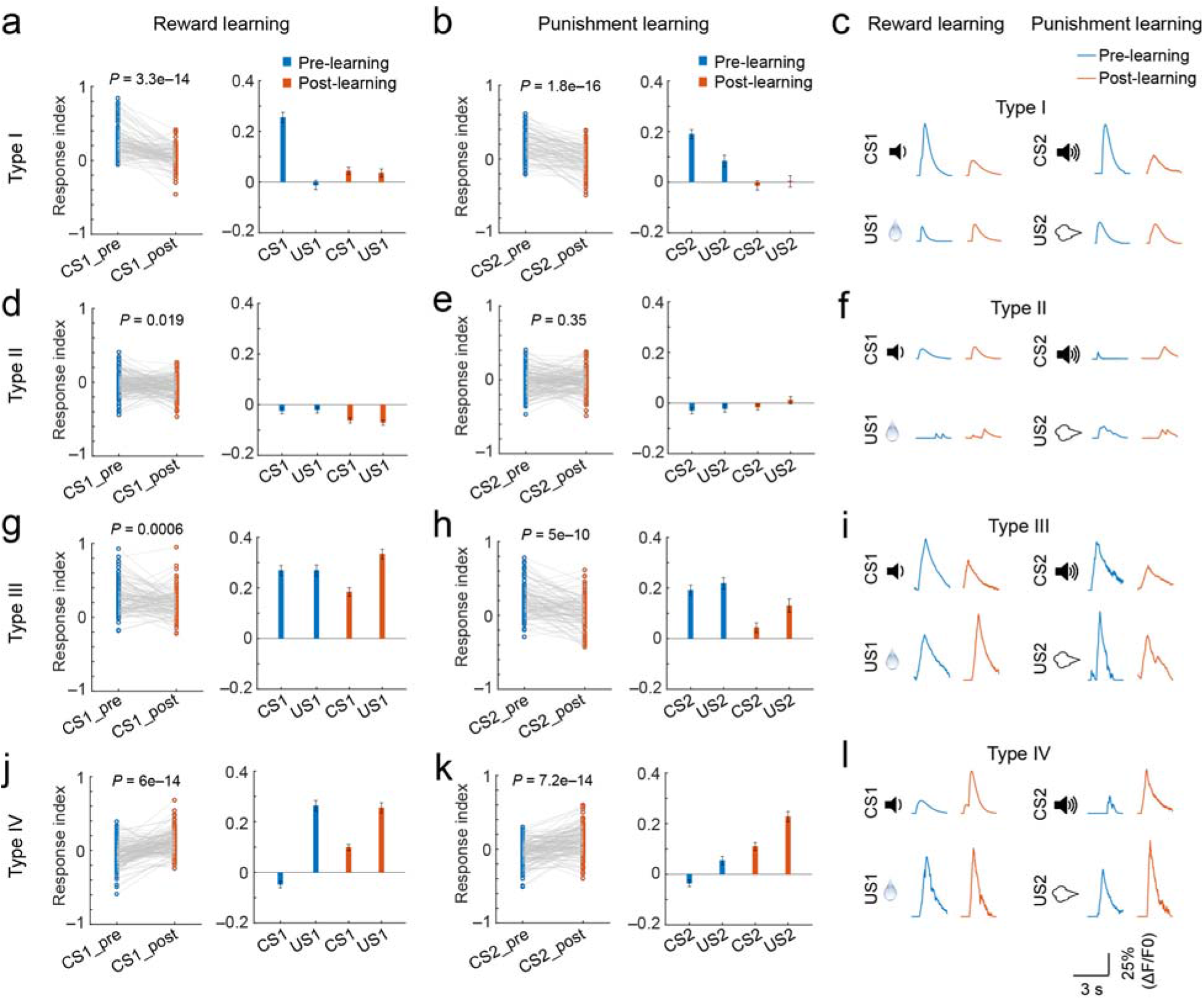
Learning induced changes in functionally distinct classes of BLA neurons. For classification of neuronal types, see Figure 5f-h. Response index is defined here as [(auROC value – 0.5) / 0.5]. (a-c) Learning induced changes in type I neurons. (a) Left: the responses of type I neurons to CS1 presentations before and after reward learning (n = 97 neurons, ***P = 3.3e–14, paired *t* test). Right: the responses of type I neurons to both CS1 and US1. The CS1 responses are the same as those shown in the left. (b) Left: the responses of type I neurons to CS2 presentations before and after punishment learning (n = 97 neurons, ***P = 1.8e–16, paired *t* test). Right: the responses of type I neurons to both CS2 and US2. The CS2 responses are the same as those shown in the left. (c) Responses of a type I neuron before and after reward learning, and those of a type I neuron before and after punishment learning. (d-f) Same as (a-c), except that the responses are from type II neurons (n = 162 neurons, *P = 0.019 in (d), P = 0.35 in (e), paired *t* test). (g-i) Same as (a-c), except that the responses are from type III neurons (n = 112 neurons, ***P = 0.0006 in (g), ***P = 5e–10 in (h), paired *t* test). (j-l) Same as (a-c), except that the responses are from type IV neurons (n = 147 neurons, ***P = 6e–14 in (j), ***P = 7.2e– 14 in (k), paired *t* test).

To further characterize how the CS responses of these BLA neurons as ensembles might transform during learning, we performed population analysis on the responses acquired before and after learning (Fig. 7) (see Methods). Specifically, we used a vector to represent the dynamic activities of each neuron in response to the presentations of CS1, CS2, US1 and US2 (Fig. 7a), during both pre-learning and post-learning periods. Thus, each cell represented one dimension in an *n*-dimensional population space (n = 518). We performed dimensionality reduction based on PCA, and used the first three principal components (PCs) to represent the ensembles of responses (to CS1, CS2, US1 and US2) before and after learning (Fig. 7b). We then computed the Mahalanobis distance between ensemble representations as a measure of similarity. Strikingly, we found that learning markedly increased the Mahalanobis distance between the ensemble representations of CS1 and CS2, but decreased that of CS1 and US1, as well as CS2 and US2. By contrast, learning did not significantly change the distance between US1 and US2 (Fig. 7b, c). These results indicate that learning increased the discriminability of CS representations, but increased the similarity between the CS and US representations; and the latter is true for both reward and punishment learning. These results are consistent with the above observations that learning caused a reduction in the fraction of neurons excited by both CS1 and CS2 (Fig. 1g, Fig. 3C), but an increase in the fraction of neurons excited (Fig. 4a) or inhibited (Fig. 4b) by both CS1 and US1, or by both CS2 and US2.

**Fig. 7.**
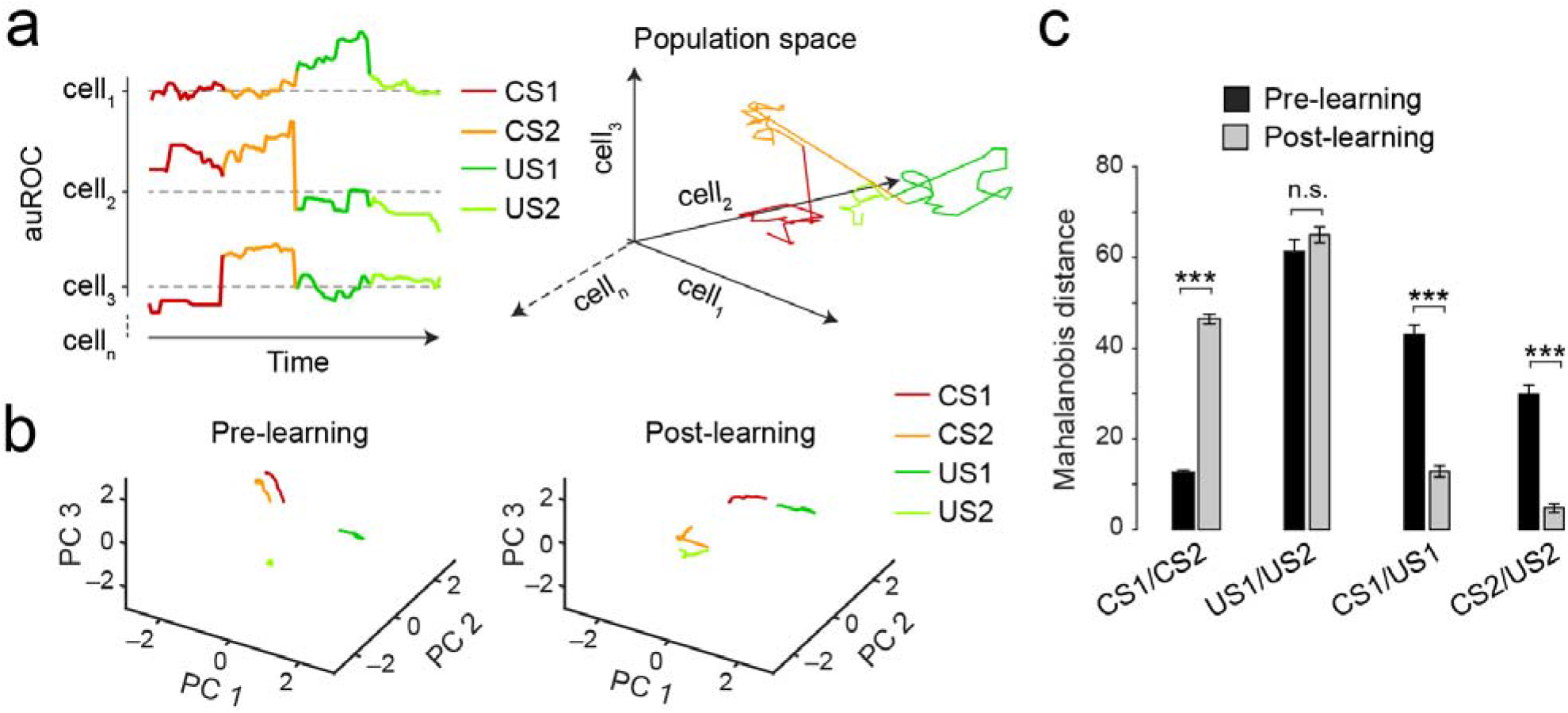
Transformation of BLA population responses during learning. (a) Left: a schematic of the population vector analysis. The dynamic activities of a neuron (cell_1_ – cell_n_) during the 3-s time window immediately after the onset of each stimulus (CS1, CS2, US1, or US2) are represented by a vector, which is composed of sequential frame-by-frame auROC values computed for that neuron. Right: the trajectory of a population vector for three example neurons in a 3D space. Each cell represents one dimension in an *n*-dimensional population space. (b) After dimensionality reduction of the population activity based on principal component analysis (PCA), the first three principal components before (left) and after (right) learning are projected onto a 3D space. (c) Quantification of the Mahalanobis distances between vectors representing neuronal responses to different stimuli (n.s., P > 0.05, ***P < 0.0001, *t* test). Data are presented as mean ± s.d.

### Reassignment of valences to CSs in the BLA during reversal learning

To examine how BLA population responses to the CSs may be updated when CS-US contingencies change, and thus guide flexible behavior, we further trained the mice (n = 6) in a reversal learning procedure, in which the initial CS-US contingencies were switched (and thus the valences predicted by the CSs reversed) without warning (Fig. 8a). We simultaneously recorded behavioral (licking and blinking) and BLA neuronal responses in trials across the reversals (Fig. 8a-e). As expected, mouse behavior changed from anticipatory blinking to anticipatory licking following the punishment-to-reward reversal (Fig. 8b), and vice versa following the reward-to-punishment reversal (Fig. 8c). Because the mice reversed their behavioral responses quickly (within 50 and 30 trials, respectively, after the punishment-to-reward and reward-to-punishment reversal trials (Fig. 8b, c), we were able to track the activities of the same population of neurons in each mouse in a single imaging session spanning all trials across the reversals (Fig. 8a, d, e).

**Fig. 8.**
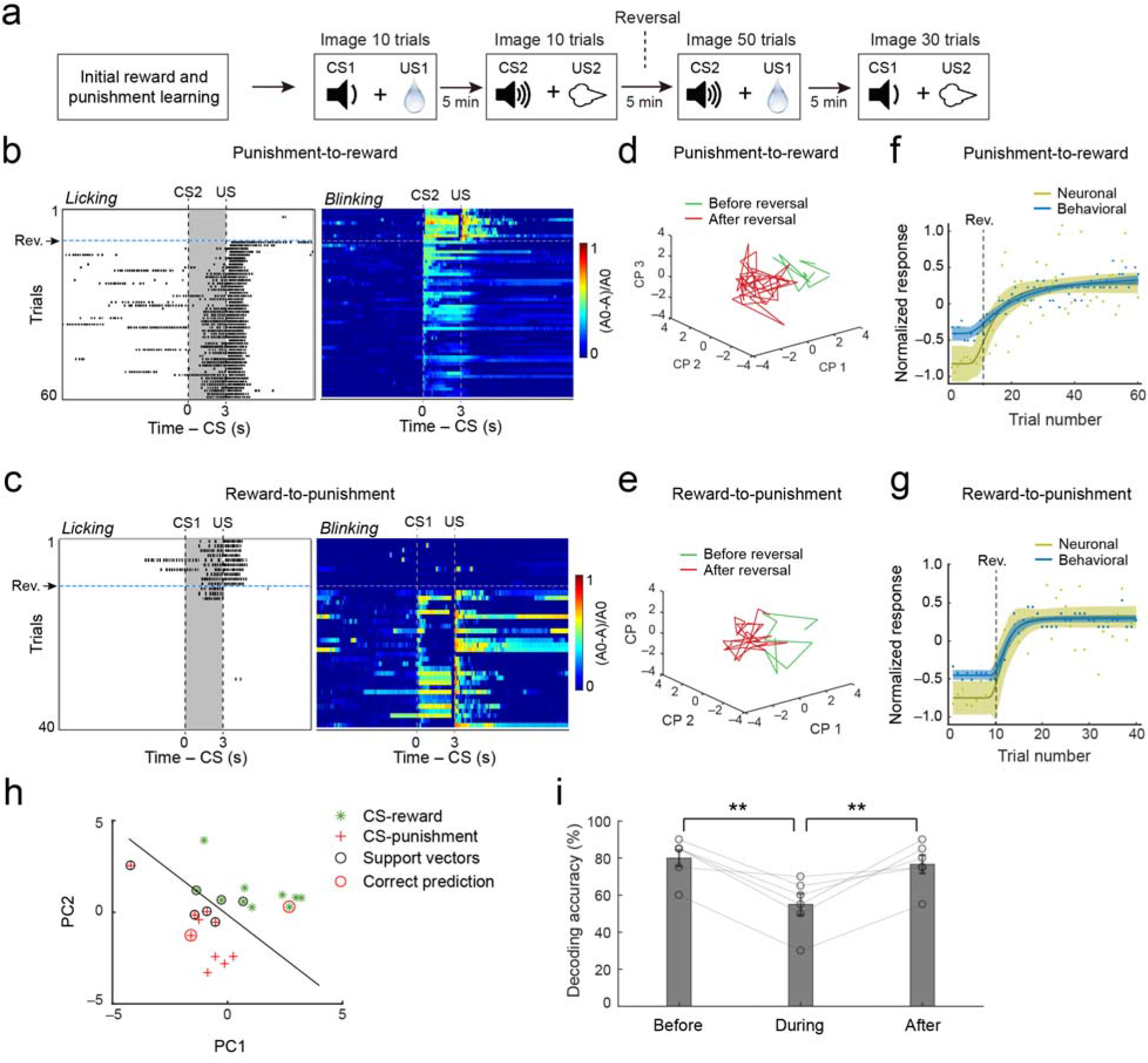
Reassignment of valences to CSs in the BLA during reversal learning. (a) A schematic showing the experimental procedure. The imaging procedure was designed such that no detachment/reattachment of the camera was needed, allowing ambiguous tracking of the same neurons throughout the reversal learning. (b) Punishment-to-reward reversal learning. Simultaneous measuring of licking (left) and blinking (right) behavior in the 10 trials before and 50 trials after the valence associated with CS2 changed from negative to positive. The horizontal dashed lines denote the first trial (11^th^ trial) at which the valence was reversed. (c) Reward-to-punishment reversal learning. Simultaneous measuring of licking (left) and blinking (right) behavior in the 10 trials before and 30 trials after the valence associated with CS1 changed from positive to negative. The horizontal dashed lines denote the first trial (11^th^ trial) at which the valence was reversed. (d, e) Projection of the trial-by-trial CS population responses (from 677 neurons) onto a 3-dimensional PCA space, in the punishment-to-reward (d) and the reward-to-punishment (e) reversal learning. (f, g) Average normalized neuronal and behavioral responses plotted as a function of trial number, for punishment-to-reward (f) and reward-to-punishment (g) reversal learning. (f) Fitting for behavioral responses, R^2^ = 0.82, fitting for neuronal responses, R^2^ = 0.60, Pearson correlation coefficient between behavioral and neuronal responses, r = 0.9756, p < 0.001. (g) Fitting for behavioral responses, R^2^ = 0.91, fitting for neuronal responses, R^2^ = 0.69, Pearson correlation coefficient between behavioral and neuronal responses, r = 0.9979, p < 0.001. Each of the dashed lines indicates the first trial (11^th^) at which the reversal of a CS valence has occurred. Shaded areas indicate 95% prediction intervals for Weibull fitting (n = 6 mice). (h) CS-reward and CS-punishment presentations are accurately classified by a linear decoder trained based on BLA population CS responses before the reversal in one mouse. (i) Performance of the decoders trained using population CS responses immediately before (“before”), immediately after (“during”), and at the end of (“after”) the reversal (n = 6 mice, before vs. during, **p = 0.0021, after vs. during, **p = 0.0082, paired t test).

To facilitate visualization of BLA population activities during the reversal learning, we projected the trial-by-trial population responses (n = 677 neurons) to presentations of the CSs onto a 3-dimensional PCA space (Fig. 8d, e). For both the punishment-to-reward (Fig. 8d) and the reward-to-punishment (Fig. 8e) reversal learning, the trajectories of the responses before the reversal are clearly separable from those after the reversal, suggesting that the CS representations were reshaped or remapped onto different neural ensembles following the switch in CS-US contingencies.

Next, we examined the trial-by-trial relationship between BLA neuronal responses and behavioral reactions before and after the valence reversals (punishment-to-reward reversal, Fig. 8f; reward-to-punishment reversal, Fig. 8g). To quantify the changes in population activity in the BLA on a trial-by-trial basis in each mouse from pre-reversal conditions, we computed the Mahalanobis distances between vectors – each representing the population CS response in a trial – and a vector distribution, which represents the population CS responses in all the 10 trials before the reversals. These distances were normalized for each mouse and subsequently averaged for each trial (Fig. 8f, g). To quantify the changes in animal behavioral responses during the reversal learning, we normalized and combined the licking and blinking responses (see Methods).

These behavioral responses were also averaged for each trial across animals. The trail-by-trail neuronal and behavioral responses were both fitted by sigmoidal Weibull functions, as previously described (Paton et al., 2006) (Methods). We found that the changes in trajectory of the BLA population CS responses were highly correlated with those of the behavioral responses in either the punishment-to-reward or the reward-to-punishment reversal learning (in both cases, Pearson correlation coefficients > 0.95, p < 0.001) (Fig. 8f, g), suggesting that the valence-specific expectation signals in the BLA at population level can be used to influence the on-going behavior.

Since the population CS responses of BLA neurons were strongly correlated with animals’ performance during reversal learning (Fig. 8f, g), we tested whether these neuronal responses can be used to predict reward or punishment deliveries. We used the population responses to presentations of both the reward-predicting CS (CS-reward) and the punishment-predicting CS (CS-punishment) in the trials before, immediately after, or at the end of the valence reversals to train linear decoders that distinguish trials in which a CS-reward was presented from those in which a CS-punishment was presented (Fig. 8h, i; Methods). We found that the decoder trained based on CS responses before or at the end of the reversals, when behavioral performance was high, had superior performance than the decoder trained based on CS responses immediately after the reversals (Fig. 8i), a time point when behavioral performance was poor (Fig. 8f, g). These results suggest that the coding fidelity in BLA neural ensembles decreases immediately after a change in CS-US contingency but recovers with relearning.

## Discussion

In this study, we imaged BLA neuronal activities in mice performing associative learning tasks driven by reward and punishment. We found that these USs of opposing valences are represented by distinct but intermingled BLA neurons. Learning reduces noise correlations in the BLA and transforms CS representations in ensembles of BLA neurons, causing the population representations of CSs paired with reward and punishment to resemble those of the actual reward and punishment, respectively, and, as a result, causing the representations of CSs to be distinct from each other. This transformation is accompanied by the emergence of prominent inhibition and plasticity in the CS responses in individual BLA neurons. Furthermore, our results suggest that reversal learning induces remapping of CS representations onto BLA ensembles representing different valences, and that this remapping may be critical for the switch in behavioral actions following a change in CS-US contingency.

The plasticity of CS response that we observed in individual BLA neurons may account, at least in part, for the learning-induced transformation of population CS representations, and for the generation of valence-specific CS responses that correlate with reversal learning. The prevalence of inhibition in BLA neurons induced by learning has not been noted before. We postulate that this inhibition may result from the learning-dependent recruitment of inhibitory interneurons in the BLA – a process that has been shown to play an important role in learning during fear conditioning (Wolff et al., 2014) – and may work in concert with the aforementioned plasticity to shape the valence-specific CS representations during learning.

Our results are consistent with previous findings that different populations of neurons in the BLA respond to punishment and reward (Belova et al., 2007; Gore et al., 2015; Kim et al., 2016; Namburi et al., 2015; Paton et al., 2006; Stuber et al., 2011), that the valence-specific CS responses of BLA neurons at single cell level correlate with valence-specific behaviors in well trained animals (Belova et al., 2007; Paton et al., 2006). Our results are also consistent with recent findings in fear conditioning, in which learning drives the population representation of CS to match that of US in the BLA (Grewe et al., 2017). Our study extends these previous studies by showing how valence-specific CS representations evolve and transform during the course of learning drive by either reward or punishment, and how reassignment of valences to sensory stimuli might occur in BLA populations and guide flexible behaviors during reversal learning.

The BLA is tightly connected to areas that are involved in the generation of behaviors motivated by negative or positive valence; it also receives inputs of all sensory modalities (Janak and Tye, 2015). Thus, BLA neurons are anatomically poised to participate in the formation of CS-US associations and contribute to the establishment of behavioral responses driven by USs of different valences. Recent studies also provide evidence suggesting that some of the BLA neurons responsive to certain kinds of reward or punishment are hard wired and can be defined either genetically (Kim et al., 2016) or by projection targets (Namburi et al., 2015; Stuber et al., 2011). An important next step is to determine the relationship between the valence-specific BLA neurons classified on the basis of their in vivo activity (Belova et al., 2007; Paton et al., 2006) (And the current study), expression of the immediately early genes or other genes (Gore et al., 2015; Kim et al., 2016; Redondo et al., 2014), and specific projection targets (Namburi et al., 2015; Stuber et al., 2011).

## AUTHOR CONTRIBUTIONS

X.Z. and B.L. conceived and designed the study. X.Z. conducted experiments and analyzed data. B.L. and X. Z. wrote the paper.

## ACKNOWLEDGMENTS

We thank Drs Pengcheng Zhou and Liam Paninski for helping with imaging data analysis, Drs. C. Daniel Salzman, Dinu F. Albeanu for critically reading an early version of the manuscript, Ga-Ram Hwang and Dylan Rebolini for technical assistance, and members of the Li laboratory for helpful discussions. This work was supported by grants from the National Institutes of Health (NIH) (R01MH101214, R01MH108924, R01NS104944, B.L.), Human Frontier Science Program (RGP0015/2016, B.L.), NARSAD (23169, B.L.), the Stanley Family Foundation (B.L.), Simons Foundation (344904, B.L.), Wodecroft Foundation (to B.L.) and the Cold Spring Harbor Laboratory and Northwell Health Affiliation (B.L.).

## METHODS

### Animals

Wild-type C57BL/6 mice (3 female, 3 male, 8–12 weeks old; The Jackson Laboratory) were used for all the experiments. Before surgery, mice were housed under a normal 12-h light/dark cycle (7 a.m. to 7 p.m. light) in groups of 2–5 animals, with food and water available *ad libitum* before behavioral training. After surgery, mice with GRIN lens implantation were housed singly. All behavioral experiments were performed during the light cycle. All animal procedures were approved and executed in accordance with Institutional Animal Care and Use Committees of Cold Spring Harbor Laboratory and with US National Institutes of Health standards.

### Viral vectors

The AAV1-Syn-GCaMP6f.WPRE.SV40 virus was purchased from the Penn Vector Core (Philadelphia, PA) and was used for expressing GCaMP6f in BLA neurons. The virus was stored in aliquots at –80 °C until use. We waited for at least 5 weeks after injection for sufficient viral expression.

### Stereotaxic surgery

Standard surgical procedures were used for stereotaxic injection and implantation, as previously described (Stephenson-Jones et al., 2016; Yu et al., 2017). Briefly, mice were anaesthetized with isoflurane (3% at the beginning and 1% for the rest of the surgical procedure), and were positioned in a stereotaxic injection frame and on top of a heating pad maintained at 35°C. A digital mouse brain atlas was linked to the injection frame to guide the identification and targeting of the amygdala (Angle Two Stereotaxic System, myNeuroLab.com). All subjects underwent two consecutive procedures in the same surgery: viral injection and GRIN lens implantation.

For each animal we made a small cranial window (1–2 mm^2^), through which a glass micropipette (tip diameter, ∽5 μm) containing the GCaMP6 virus (1:4 or 1:8 dilution) was lowered down to the target. Virus (∽0.3 μl) was delivered with pressure applications (5–20 psi, 5–20 ms at 0.5 Hz) controlled by a Picrospritzer III (General Valve) and a pulse generator (Agilent). The injection was performed using the following stereotaxic coordinates for the BLA: −1.6 mm from Bregma, 3.3 mm lateral from midline, and 4.5 mm vertical from cortical surface. The speed of injection was ∽0.1 μl/10 min.

After virus injection, we waited for at least 10 min before removing the injection pipette. A GRIN lens (diameter, 0.6 mm; length, 6.7 mm; Inscopix) was then carefully implanted 200 μm above the center of the injection using a GRIN lens holder (Inscopix). The speed for lowering the GRIN lens was constant and slow (∽100 μm/min). We secured the GRIN lens to the skull with C&B-Metabond Quick adhesive luting cement (Parkell Prod), and subsequently mounted a small piece of metal bar on the skull for head-fixation. Four to six weeks following GRIN lens implantation, we checked the fluorescent signals using a miniature microscope (nVista HD, Inscopix) in these mice under awake and head-fixation conditions. A baseplate (Inscopix) attached to the miniature microscope was then positioned above the GRIN lens. The focal plane was adjusted slowly until vascular structures and GCaMP6 dynamic activities were clearly observed (Resendez et al., 2016). The baseplate was subsequently secured with dental cement.

### Immunohistochemistry

Immunohistochemistry experiments were performed following standard procedures. Briefly, mice were anesthetized with Euthasol (0.2 ml; Virbac, Fort Worth, Texas, USA) and transcardially perfused with 30 ml of PBS, followed by 30 ml of 4% paraformaldehyde in PBS. Brains were extracted and further fixed in 4% PFA overnight followed by cryoprotection in a 30% PBS-buffered sucrose solution for 36 h at 4 °C. Coronal sections (40 or 50 μm thickness) were cut using a freezing microtome (Leica SM 2010R, Leica). Sections were first washed in PBS (3 × 5 min), incubated in PBST (0.3% Triton X-100 in PBS) for 30 min at room temperature (RT) and then washed with PBS (3 × 5 min). Next, sections were blocked in 5% normal goat serum in PBST for 30 min at RT and then incubated with the primary antibody overnight at 4 °C. Sections were washed with PBS (5 × 15 min) and incubated with the fluorescent secondary antibody at RT for 2 h. After washing with PBS (5 × 15 min), sections were mounted onto slides with Fluoromount-G (eBioscience, San Diego, California, USA). Images were taken using a LSM 780 laser-scanning confocal microscope (Carl Zeiss, Oberkochen, Germany). The primary antibody used was rabbit anti-GABA (Sigma, St. Louis, MO, USA; catalogue number A2052). The fluorophore-conjugated secondary antibody used was Alexa Fluor^®^ 594 donkey anti-rabbit IgG (H+L) (Life Technologies, Carlsbad, California, USA; catalogue number A21207).

### Behavioral training

Water deprivation started 23 hours before training in an auditory classical conditioning task, during which mice were head restrained using custom-made clamps and metal head-bars. Each mouse was first habituated to head restraint for 2–3 days prior to training. Unpredicted drops of water (5 µl) were delivered during the habituation. Once animals have learned how to lick, they were subjected to conditioning wherein two distinct auditory cues (conditioned stimuli, CS) were associated with different outcomes (unconditioned stimulus, US): a 2-kHz tone (1 s) predicted that a water reward (5 µl) was available from a metal spout next to the mouth, whereas a 10-kHz tone (1 s) predicted that an unpleasant air puff (40 psi, 100 ms) would be blown to the face, in an area close to the eye. Animals were trained one session per day, with each session consisting of a reward block (100 trials) followed by a punishment block (30–50 trials). Each trial began with a CS, followed by a 2 second delay, and then a US. The inter-trial interval was randomly variable between 40–50 s.

Once mice have learned the initial associations (with the criterion that they correctly predicted the outcomes in more than 90% of the trials), they were further trained in a reversal learning session, in which we reversed the CS-US contingencies such that the CS initially associated with punishment became associated with reward (the punishment-to-reward reversal), and the CS initially associated with reward became associated with punishment (the reward-to-punishment reversal). Specifically, we first “reminded” mice with the original CS-US contingencies (10 trials of CS1-reward pairing, followed by 10 trials of CS2-punishment pairing). We then immediately subjected these mice to two blocks of reversal training, with each reversal being initiated without warning (50 trials of CS2-reward paring, followed by 30 trials of CS1-punishment pairing).

### Behavioral data collection and analysis

A custom software written in LabView (National Instruments) was used to control the delivery of CS and US and record behavioral responses, including licking and eye-blinking, during the CS-reward and CS-punishment conditioning. A metal spout was placed in front of animal mouth for water delivery. The spout also served as part of a custom “lickometer” circuit, which registered a lick event each time a mouse completed the circuit by licking the spout. The lick events were recorded by a computer through the LabView software.

Eye blinking was tracked using a high-speed camera (FL3-U3-13S2C-CS, 120 HZ, Point Grey), which was controlled by a Bonsai software (Bonsai). Offline video analysis was conducted using EthoVision XT software (Noldus; Wageningen, The Netherlands). To measure the size of the eye, we manually selected a region of interest (ROI) surrounding the eye. Pixels corresponding to the eye were assigned as those that were darker than the surrounding background within the ROI. To quantify the changes in eye size (ΔA, which we refer to as “blinking”, although mice tend to close their eyes in response to an airpuff for a period much longer than that of a typical blinking event), we computed ΔA/A0(t) = (A(t) – A0)/A0, where A0 is the median size of the area corresponding to the eye during the 10-s baseline before CS onset, using a custom script written in MATLAB (The MathWorks, Inc., Natick, Massachusetts, USA). A blinking event was defined as ΔA/A0(t) < 20%.

### *In vivo* calcium imaging data acquisition and analysis

We followed a recently described procedure for the *in vivo* imaging experiments (Resendez et al., 2016; Yu et al., 2017). All imaging experiments were conducted on awake behaving mice under head-restraint in a dark, sound attenuated box. GCaMP6f fluorescence signals were acquired using a miniature integrated fluorescence microscope system (Inscopix, Palo Alto, CA) through GRIN lenses implanted in the BLA.

We installed a baseplate on top of the GRIN lens for each mouse, as described previously (Resendez et al., 2016; Yu et al., 2017). Before each imaging session, the miniature microscope was attached to the baseplate. The analog gain (1–3) and LED output power (10–40% of the maximum; 0.1–0.4 mW) of the microscope were set to be constant for the same subject across imaging sessions. The microscope was adjusted such that the best dynamic fluorescence signals were at the focal plane, which was subsequently kept constant across imaging sessions. To synchronize sensory stimuli and behavioral events with imaging acquisition, the Data Acquisition Box of the nVista Imaging System (Inscopix, Palo Alto, CA) was triggered by a behavioral control software written in LabView (National Instruments) through an NI data acquisition device (USB6008, National Instruments, CA). Compressed gray scale images were then recorded with nVistaHDV2 (Inscopix) at 10 frames per second. During imaging, the time stamps of different events, including the trigger signals sent to the microscope, CS, US, licks, and blinks, were all recorded with the behavioral control software (written in LabView) running on a dedicated high-speed computer.

We imaged BLA neuron activities during pre-learning, post-learning, and reversal learning sessions. To reliably detect stimulus-driven responses while minimizing photobleaching, we typically imaged neuronal responses to the same stimulus in 10 trails (except for the reversal learning; see below), with the imaging duration for each trial being 23 s to cover baseline, CS and/or US responses. During habituation, we imaged the responses to either CSs or USs, which were presented randomly interleaved.

During conditioning, 3 mice were able to learn both the CS-reward and the CS-punishment associations within the first session. We were thus able to image the same population of neurons in each of these mice before and after learning (Fig. 5). For each of the other 3 mice, we carefully adjusted the camera to the similar focal plane as that in the habituation session (Fig. 1), and then imaged the CS1-US1 and the CS2-US2 associations once the animal reached a high successful level (90%). Note that we did not attempt to track the same neurons during imaging in these other 3 mice that took more than one session to learn the tasks (Fig. 1 & 3), because imaging in these mice required disassembling and reassembling the camera across sessions or days, which inevitably caused changes in the focal plane and thus the identities of neurons in the field of view.

For reversal learning, we first acquired imaging data under the original CS-US contingencies (10 trials of CS1-reward pairing, followed by 10 trials of CS2-punishment pairing). This was immediately followed by imaging throughout the reversal learning (50 trials of CS2-reward paring, and then 30 trials of CS1-punishment pairing).

For imaging data processing and analysis, we first used Mosaic (version 1.0.0b; Inscopix, Palo Alto, CA) to combine all the video clips, each of which was recorded from one imaging session, into a single image stack (in TIFF format). The image stack was then spatially down sampled by a factor of 4 and corrected for motion artifacts using Mosaic. The motion-corrected video was next cropped to delete the margin areas.

Next, to address the problem of high levels of background fluorescence intrinsic to one-photon imaging, we applied the newly developed image analysis method, extended constrained nonnegative matrix factorization (CNMF-E) (Yu et al., 2017; Zhou et al., 2018), which models the background with two realistic components: the constant baseline of each pixel, and the fluctuations from out-of-focus signals that is constrained to have low spatial-frequency structure. This decomposition avoids cellular signals being absorbed into the background term. After subtracting the background approximated with this model, we used constrained non-negative matrix factorization (CNMF) to demix neural signals and get their denoised and deconvolved temporal activity, termed ΔF (Pnevmatikakis et al., 2016). The CNMF-E method was carried out using a custom Matlab algorithm (for a detailed description of this method, see (Zhou et al., 2018)). We then normalized ΔF by F0 to get ΔF/F0, where F0 is the modeled background fluorescence intensity.

Once the temporal activity of the neurons was extracted, we characterized the CS and US responses of each neuron using ROC (receiver-operating characteristic) analysis, in which we compared the median ΔF/F0 values during the baseline periods (10 s immediately prior to the delivery of CS or US) and those during CS or US presentations (3 s immediately after the onset of CS or US) in all trails with a criterion moving from zero to the maximum ΔF/F0 value. We then plotted the probabilities that the ΔF/F0 values during CS or US presentations were greater than the criteria against the probabilities that the baseline ΔF/F0 values were greater than the corresponding criteria. The area under the ROC curve (auROC) quantifies the degree of overlap between the two probability distributions (i.e., the discriminability of the two). The values of auROC vary between 0 and 1, with values greater than 0.5 indicating increased activity during CS or US presentations relative to during baseline, and values less than 0.5 indicating decreased activity during CS or US presentations relative to during baseline. A permutation test (iteration 5000 times) was used to determine whether the median ΔF/F0 values during CS or US presentations were significantly higher or lower (P < 0.05) than during baseline, and thus classify a neuron as being excited or inhibited, respectively, by CS or US.

### Classification of neurons with clustering analysis

Briefly, to classify neurons based on their CS and US responses before and after learning (Fig. 5f-h), the ROC curve was generated as described above in the “*In vivo* calcium imaging data acquisition and analysis” section. We performed principal component analysis (PCA) on these auROC values for all cells (Cunningham and Yu, 2014). We subsequently applied hierarchical clustering analysis to the first three principal components (PCs) using a correlation distance metric and complete agglomeration methods (Stephenson-Jones et al., 2016).

### Analysis of noise correlations

Noise was defined as the trial-to-trial fluctuations around the mean in responses to repeated presentations of CS1 or CS2. Noise correlation was quantified as the Pearson correlation coefficient between such fluctuations in a pair of neurons. For this analysis, we used an auROC value to represent the CS response of a neuron in each trial, which was computed based on activities in the 3-s time window immediately before and that immediately after the CS onset.

### Population vector analysis

To investigate the learning-induced changes in the responses to CS or US at population level, we performed population vector analysis, adapted based on that described in a recent study (Rozeske et al., 2018). Briefly, we created a series of *n*-dimensional (*n* equals the number of neurons) activity vectors by pooling the auROC values of individual neurons at each time point. Therefore, the ensemble BLA response at a particular time point is represented by a vector with a dimension equal to the total number of neurons in that ensemble. We used PCA for dimensionality reduction, and projected the population vectors onto a three-dimensional space for data visualization. To examine whether learning induced changes in BLA population responses to CS and US, we computed the Mahalanobis distances between vectors. For example, the Mahalanobis distance (MD) between responses to CS1 and CS2 at each time point is defined by: 
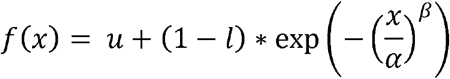
where PV(CS1) and PV(CS2) are the population vectors of responses to CS1 and CS2, respectively. S^−1^ is the inverse of the covariance matrix.

### Correlation analysis of neural and behavioral responses

To compare the trajectory of changes in behavioral responses with that in neural responses, we averaged the normalized behavioral (licking and blinking) and neuronal responses for each animal in the 10 trials prior to and 50 (punishment-to-reward reversal) or 30 (reward-to-punishment reversal) trials after the valence reversal, as previously described (Paton et al., 2006). In brief, we normalized the trial-by-trial behavioral responses by dividing them with the median response, followed by subtracting from the resulting responses the mean of the normalized values. To make the behavioral responses go from low to high, we multiplied the values by –1 for responses (licking or blinking) that were higher before than after the reversal. For neuronal responses, we first performed PCA on the trial-by-trial CS responses (using auROC values) of all neurons in each mouse across reversal learning, and used the first three components to represent the population CS responses. We then computed the Mahalanobis distance between the vector representing the population CS response in each trial and a vector distribution representing the population CS responses in all the 10 trials before the reversal. We normalized these distances, which represent the changes in CS responses during reversal learning, using a procedure similar to that used for normalization of the behavioral response. We next fitted both behavioral and neural data with sigmoidal Weibull functions (Paton et al., 2006):

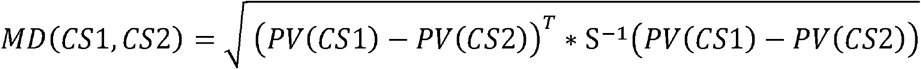

where x is the behavioral or neural responses, u and l set the upper and lower asymptotes, respectively, and α and β determine the latency and abruptness, respectively, of the rise of the function. The Weibull functions model the averaged and normalized responses as a function of trial number.

To determine the relationship between neuronal responses and behavioral responses, we computed the Pearson correlation coefficients between the trial-by-trial neuronal responses and behavioral responses (licking or blinking) to the CS presentations.

### Decoding analysis

We performed the decoding analysis using the support vector machine (SVM) in MATLAB (MathWorks) to determine whether CS-reward and CS-punishment presentations could be predicted on the basis of BLA population CS responses acquired in each mouse at different stages of the reversal learning. We first conducted ROC analysis (based on activities the 3-s time window immediately before and that immediately after the CS onset) and used auROC values to represent the trial-by-trial responses of each BLA neuron to presentations of either CS-reward or CS-punishment throughout the reversal learning. For each of the 6 mice, we captured the trial-by-trial population CS responses by a vector containing the responses of all neurons from that mouse, with each neuron representing one dimension in a multidimensional population space. We applied PCA on the multidimensional data from each mouse and used the first two principal components to represent the population response in each trial. We then used the low dimensional trial-by-trial population responses from each mouse to train and test binary linear classifiers with SVM to distinguish CS-reward presentations from CS-punishment presentations. Specifically, we used the responses from the 20 trials (10 CS-reward trials and 10 CS-punishment trials) immediately before, immediately after, or at the end of the valence reversals to train and test a classifier. For decoder training and testing, we used a 10-fold cross-validation procedure, in which all datasets were randomly partitioned to 10 equal size subsamples, with each subsample containing equal number of responses from a given class (i.e., CS-reward vs. CS-punishment). We used 9 of the 10 subsamples for training and the remaining 1 for validating the decoder, and repeated this process 10 times. Thus, each of the 10 subsamples was used exactly once as the validation data. The percentage of accurate classification incidence in the 10 times was reported as the final classification accuracy.

### Statistics and data presentation

All statistics are indicated where used. Statistic analyses were performed with Matlab (MathWorks, Natick, MA). All behavioral experiments were controlled by computer systems, and data were collected and analyzed in an automated and unbiased way. Virus-injected animals in which the injection site was incorrect were excluded. If the tract and tip of the GRIN lens was outside of the targeted area, the mouse was also excluded. No other mice or data points were excluded.

**Supplementary Figure 1.**
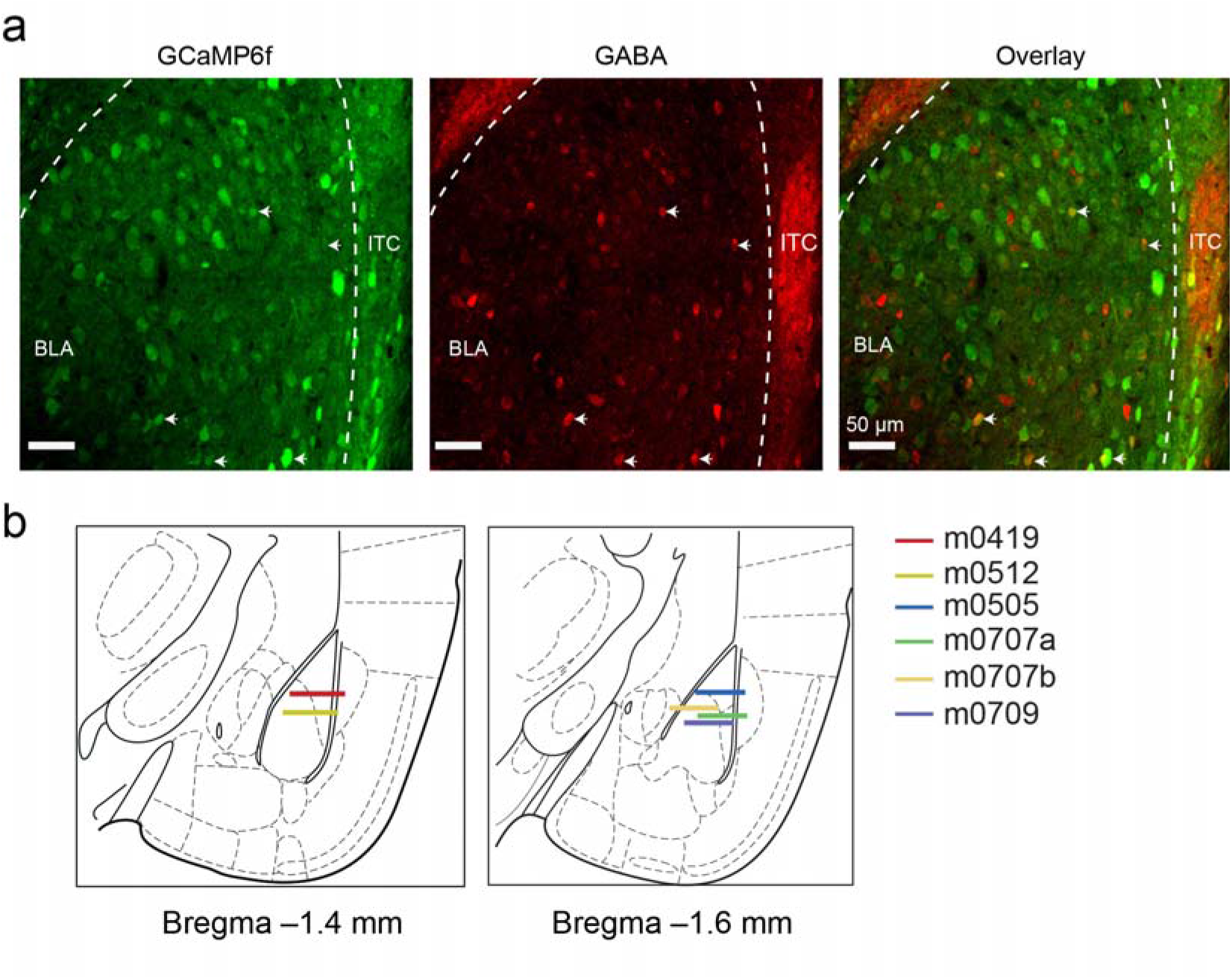
Histology on virus infection and diagrams of GRIN lens implantation. **(a)** The GCaMP6f-expressing BLA neurons were predominantly non-GABAergic. Shown are confocal images of the BLA infected with the AAV expressing GCaMP6f. GABAergic neurons were identified with an antibody recognizing GABA. White arrows indicate few cells that expressed both GCaMP6f and GABA (4.9±1% of GCaMP6f-expressing cells expressed GABA; n = 3 mice). (b) Schematics showing the positioning of the tips of GRIN lenses implanted in the BLA of all the mice (n = 6) used in the imaging experiments. Diagrams were modified from The Mouse Brain in Stereotaxic Coordinates (Compact 3^rd^ Edition, Franklin and Paxinos, 2008).

**Supplementary Figure 2.**
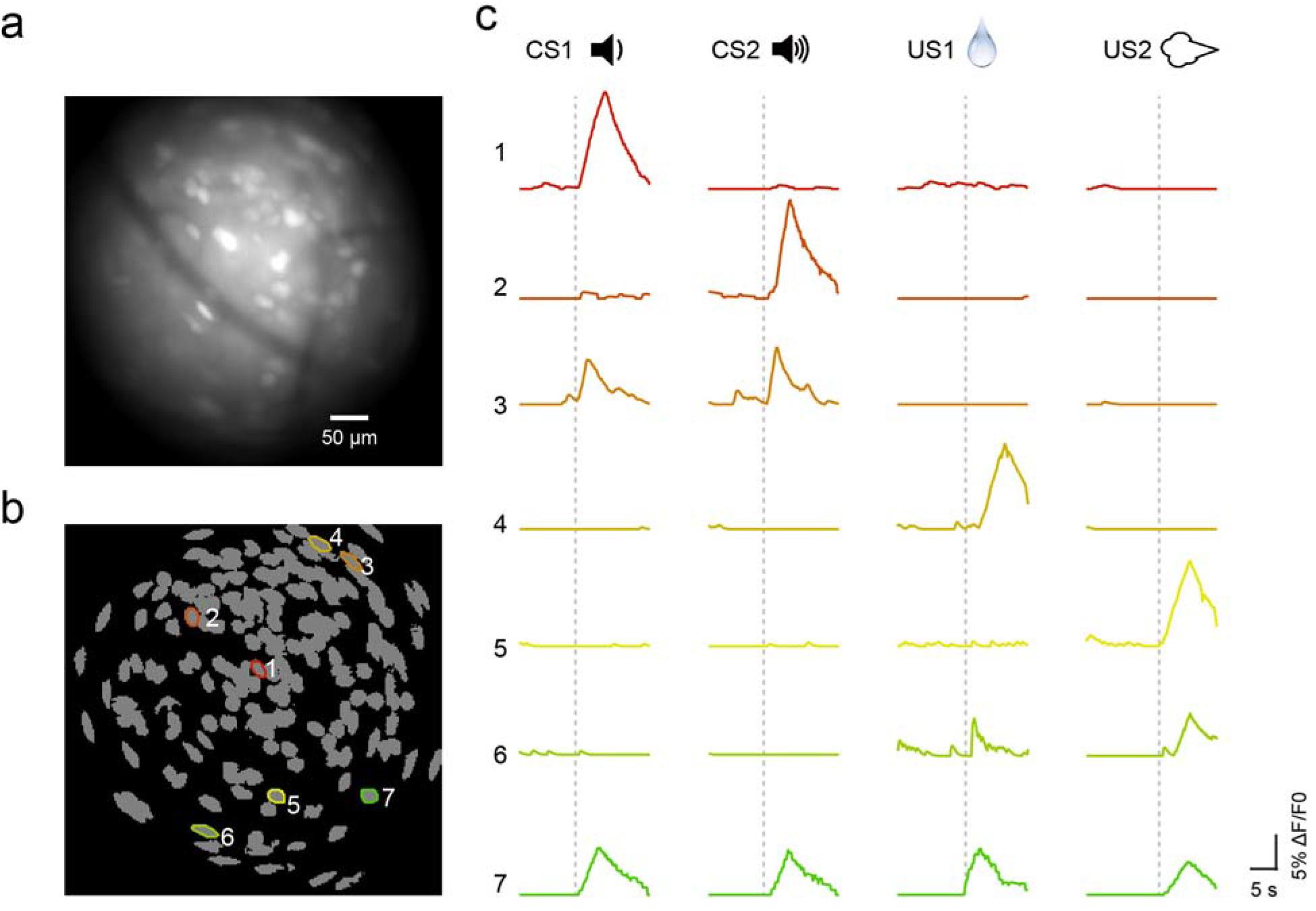
Imaging BLA neuronal activities through GRIN lenses in behaving mice. (a) The field of view through a GRIN lens implanted in the BLA, showing the raw GCaMP6f fluorescence signals from BLA neurons acquired with the miniature microscope. (b) The spatial locations of individual extracted neurons (see Methods) in the field of view shown in (a). The contours of 7 representative neurons were outlined, color-coded and numbered. (c) The temporal calcium activities of the 7 neurons outlined in (b), color-coded and numbered in the same way. Each trace represents neuronal activities recorded from a single trial. Dashed lines indicate the onset of CS or US presentations.

**Supplementary Figure 3.**
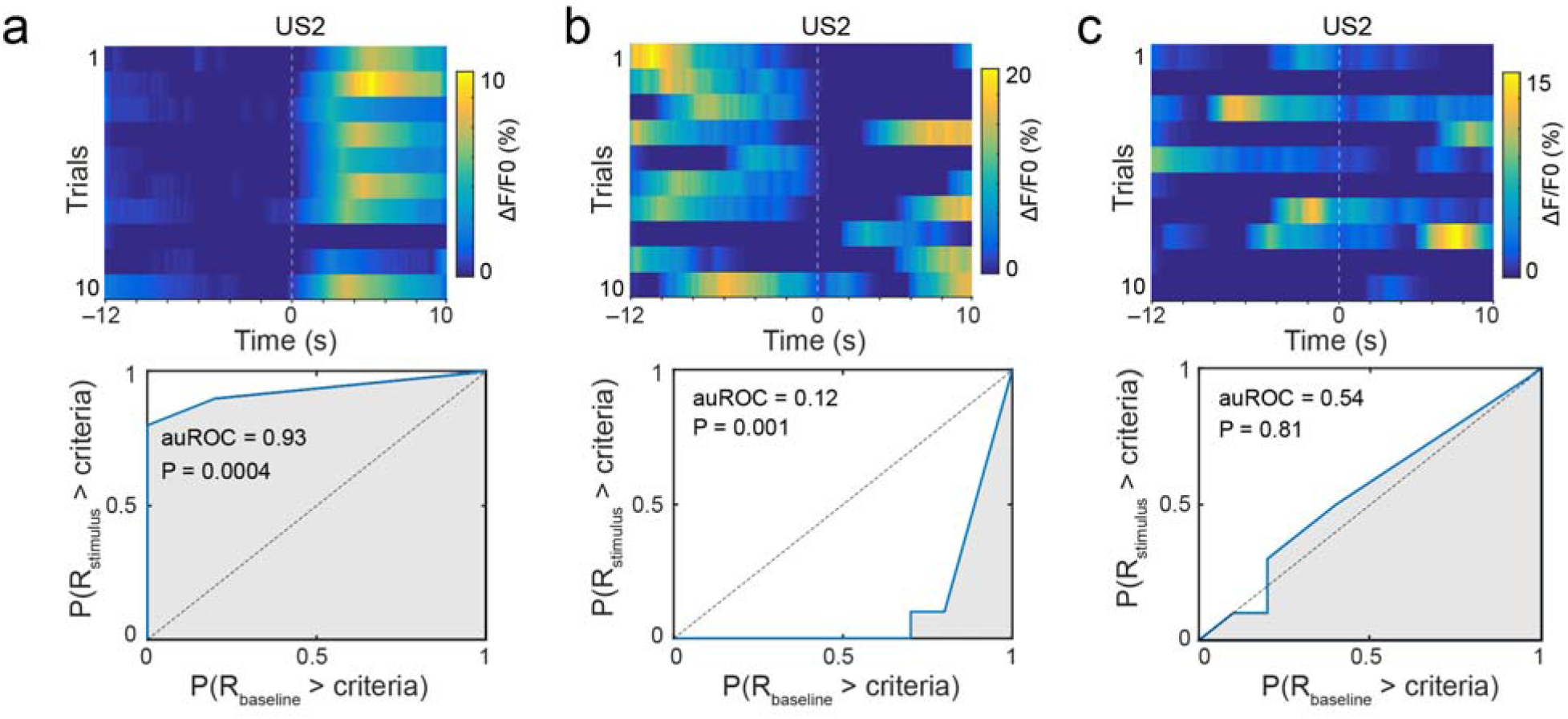
Definition of stimulus-responsive neurons. Heatmaps of trial-by-trial temporal calcium activities in three representative neurons that showed excitatory (a), inhibitory (b), or no (c) responses to US2 (airpuff). Dashed lines indicate the onsets of US2 (t = 0). Each of the bottom panels is an auROC plot for the corresponding neuron on the top panel. P values were generated from permutation tests.

**Supplementary Figure 4.**
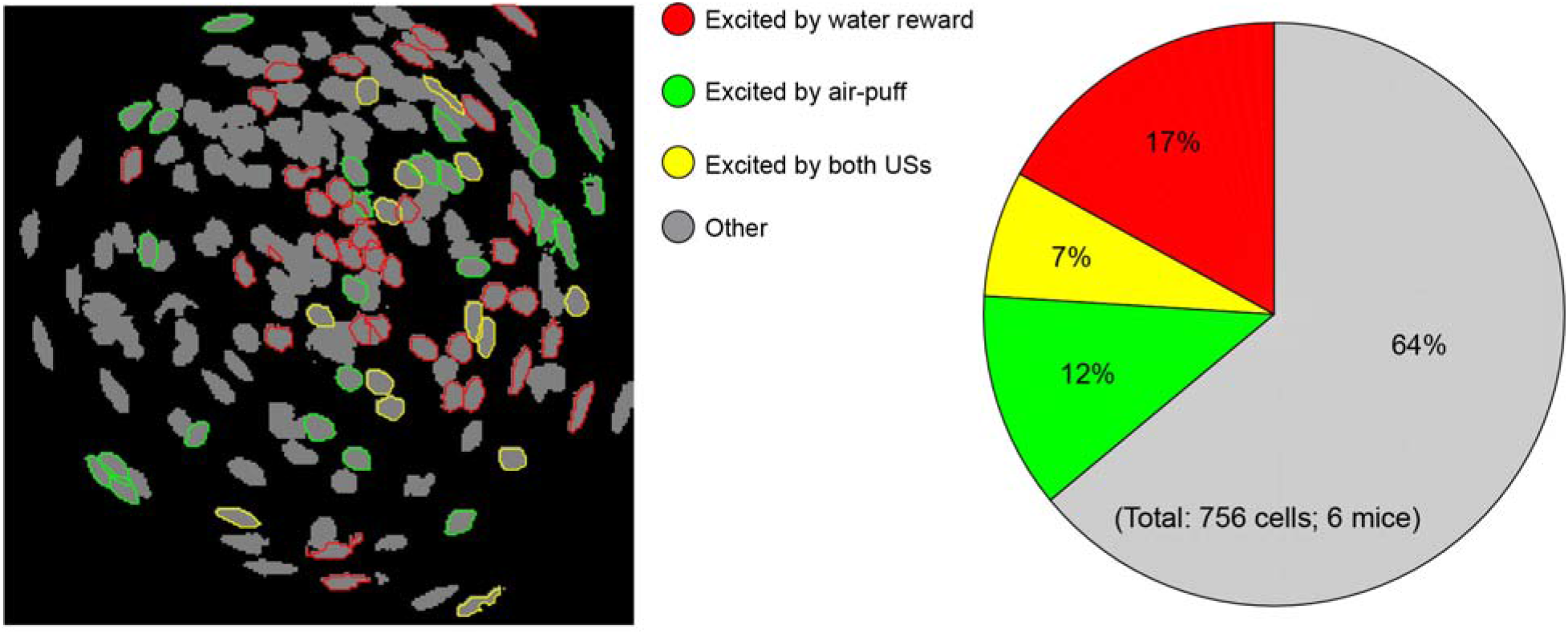
BLA neurons responsive to appetitive or aversive stimulus are spatially intermingled. Left: the spatial locations of individual extracted neurons (see Methods) in the field of view in the BLA of a representative mouse. The contours of neurons that showed significant responses to water reward, air-puff, or both stimuli were color-coded. The rest of the neurons showed either inhibitory responses or no significant response to the two stimuli. The spatial distribution of these populations does not form obvious patterns. Right: pie graph showing the percentage distribution of these BLA populations (n = 756 cells from 6 mice).

**Supplementary Figure 5.**
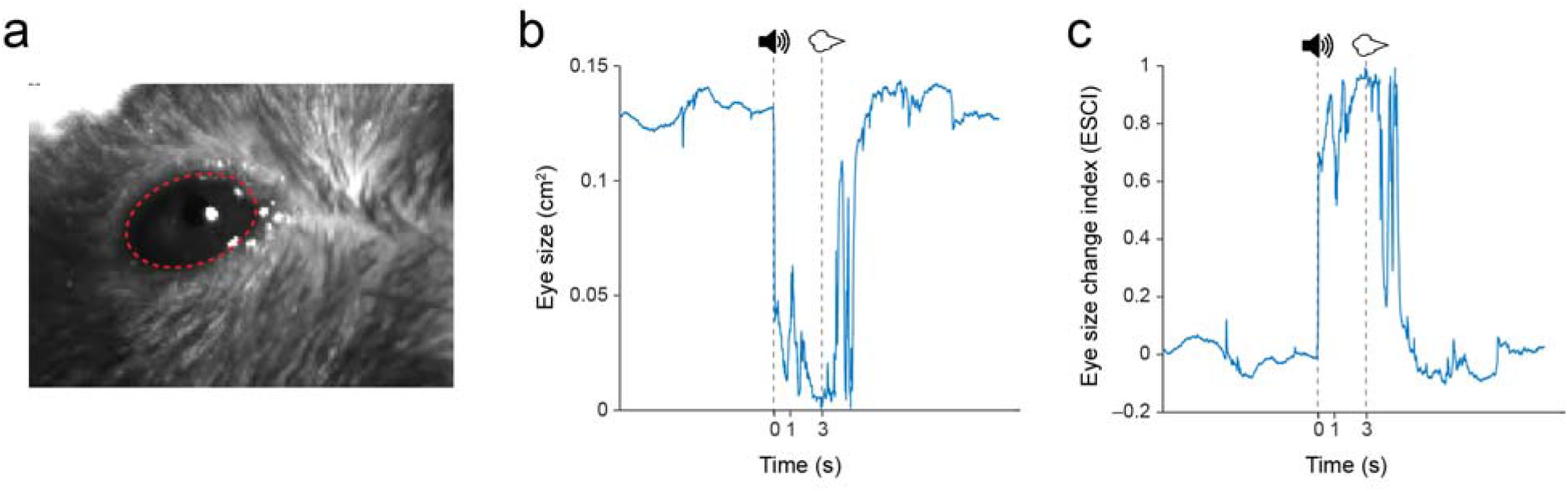
Tracking the changes in eye size. (a) The eyes of mice were tracked using a high-speed camera triggered by the behavioral control software, such that the tracking was synchronized with other events, including recording licking behavior or imaging neuronal activities. The size of the eye (outlined by the red circle) was measured offline. (b) Tracking of eye size during the punishment task. Dashed lines indicate the onsets of CS (1 s in duration) and US (air-puff) presentations. (c) The relative eye size change over time for each mouse was represented by “eye size change index (ESCI)” (ESCI = (A0–A)/A0, where A0 is the median eye size during the 10-s baseline before CS onset, and A is the eye size at a given time).

**Supplementary Video 1. Extraction of the spatial and temporal components of neuronal activity from GCaMP6 signals in BLA neurons imaged through GRIN lenses**. Shown are videos of GCaMP6f signals from BLA neurons in a representative mouse receiving air-puffs. These videos are short clips of movies stitched together, with each clip representing a trial (the onset of which is denoted by each of the solid black lines in the bottom right panel). These videos are played in synchrony, and represent the following contents: “**raw data**”, GCaMP6 signals without any processing; “**background**”, the background components in the raw data approximated with our model; “**raw–background**”, the remaining signals after subtraction of the background from the raw data; “**residual**”, the remaining signals; “**denoised**”, the denoised and deconvolved spatiotemporal activity for each neuron obtained from the “raw–background” signals using the CNMF-E (Methods). In the “denoised” video, the contours of 3 representative neurons are traced and numbered. The temporal activities of these 3 neurons are displayed in the bottom right panel, in which the moving line indicates the passage of time synchronized with all the videos; the stationary solid black lines denote the onsets of trials (and thus the junctions between the short movie clips); and the dashed red lines denote the onsets of air-puffs. Neurons #1 & 2 are air-puff-responsive neurons, whereas neurons #3 only shows spontaneous activities. The scale bar beside each of the videos denotes ΔF values. See Methods for a more detailed description.

**Supplementary Video 2. Licking behavior driven by the reward cue**. This video shows the licking behavior of a thirsty mouse after presentation of a sound (delivered at 10 s) predicting water reward (delivered at 13 s).

**Supplementary Video 3. Blinking behavior driven by the punishment cue**. This video shows the blinking behavior of a mouse after presentation of a sound (delivered at 10 s) predicting air-puff (delivered at 13 s).

